# Impact of four common hydrogels on amyloid-β (Aβ) aggregation and cytotoxicity: Implications for 3D models of Alzheimer’s disease

**DOI:** 10.1101/711770

**Authors:** Laura W. Simpson, Gregory L. Szeto, Hacene Boukari, Theresa A. Good, Jennie B. Leach

**Affiliations:** Department of Chemical, Biochemical and Environmental Engineering, University of Maryland Baltimore County, Baltimore, Maryland, USA; Marlene and Stewart Greenebaum Comprehensive Cancer Center, University of Maryland, Baltimore, Maryland, USA; Division of Physical and Computational Sciences, Delaware State University, Dover, Delaware, USA; Division of Molecular and Cellular Biosciences, National Science Foundation, Alexandria, Virginia, USA

**Keywords:** Alzheimer’s disease, beta amyloid, protein aggregation, hydrogels, confinement, cytotoxicity

## Abstract

The properties of a hydrogel utilized in 3D culture can influence cell phenotype and morphology, yielding striking similarities to cellular processes that occur *in vivo*. Indeed, research areas including regenerative medicine, tissue engineering, cancer models, and stem cell cultures have readily utilized 3D biomaterials to investigate cell biological questions. However, cells are only one component of this milieu. Macromolecules play roles as bioactive factors and physical structures. Yet, investigations of macromolecular biophysics largely focus on pure molecules in dilute solution. Biophysical processes such as protein aggregation underlie diseases including Alzheimer’s disease, which is hallmarked by accumulated neurotoxic amyloid-β (Aβ) aggregates. Previously, we demonstrated that Aβ cytotoxicity is attenuated when cells are cultured within type I collagen hydrogels vs. on 2D substrates. Here, we investigated whether this phenomenon is conserved when Aβ is confined within hydrogels of varying physiochemical properties, notably mesh size and bioactivity. We investigated Aβ structure and aggregation kinetics in solution and in hydrogels (collagen, agarose, hyaluronic acid and polyethylene glycol) using fluorescence correlation spectroscopy and thioflavin T assays. Our results reveal that all hydrogels tested were associated with Aβ cytotoxicity attenuation. We suggest that confinement itself imparts a profound effect, possibly by stabilizing Aβ structures and shifting the aggregate equilibrium toward larger species. It is likely that the milieu that exist within cells and tissues also influences protein-protein interactions; thus, we suggest that it is critical to evaluate whether protein structure, function, and stability are altered in 3D systems vs. ideal solutions and 2D culture.

## Introduction

Alzheimer’s disease (AD) is the most common form of dementia [1], and is associated with the accumulation of amyloid-β (Aβ), a protein whose aggregation is associated with neurotoxicity [2]. There is still debate over the exact size and structure of the most toxic Aβ species, but it is widely held that small oligomers that lack β-sheet structure are more toxic than assembled β-sheet fibrils [3–7]. Since the first genetic connection between Aβ and early-onset AD, investigators have targeted Aβ as a potential therapeutic strategy [8–10]. Many anti-Aβ antibody drugs (e.g., aducanumab, solanezumab, crenezumab, gantenerumab) have had promising preclinical results; however, all have failed to show a significant clinical benefit [11–13].

We have previously demonstrated that Aβ cytotoxicity was attenuated in 3D type I collagen hydrogels as compared to in 2D culture in which significant cell death occurred [14]. We suggested that in collagen hydrogels, a) the structural equilibrium of Aβ is shifted to favor larger β-sheet aggregates in contrast to in solution where the smaller oligomeric Aβ species persisted and b) that this shift in distribution of Aβ structures may have led to the stabilization of larger, less toxic fibril species compared to the species observed in solution. Confinement excludes the locally-available solvent, which promotes a more compact peptide/protein structure. Confinement also increases local protein concentration, promoting protein-protein interactions. This finding challenges the choice of 2D culture for investigations of Aβ cytotoxicity. Yet, only a few 3D gel-based models of AD have been published to date, all using the gel matrix Matrigel® (Corning) [15–17]. Matrigel® is composed of basement membrane extracellular matrix (ECM) molecules (60% laminin, 30% collagen IV, and 8% entactin) and is also commonly used to investigate stem cell differentiation [18–22].

A second possible explanation of our previous results is that 3D culture in a collagen hydrogel results in changes in cell signaling, phenotype, or potentially the expression or function of receptors available for Aβ interaction, resulting in attenuated toxicity. In support of this explanation, it is known that epigenetic changes occur in 3D culture that influence cellular phenotype [23–24]. Further, in comparison to 2D culture, cell morphologies of neuronal cells grown in 3D culture are strikingly similar to those expressed *in vivo* [25–29]. Finally, there have been numerous reports of cell surface receptors that bind Aβ, with the numbers of candidate receptors totaling 30 or more [30]. Thus, it is possible that the attenuation of Aβ cytotoxicity observed in 3D collagen may be unrelated to Aβ structural changes, but instead, be related to cellular responses that are altered due to 3D culture or the presence of collagen.

In this work, we investigated the extent to which of this Aβ confinement effect also occurs in other 3D hydrogels that vary in biomaterial physiochemical properties (e.g., mesh size, chemical composition, and biological activity). In this work, we studied Aβ structure, aggregation, and toxicity in hydrogels primarily composed of type I collagen, low melting temperature agarose, hyaluronic acid (HA), or polyethylene glycol (PEG) (Figure 1). In choosing these gel types, we were less concerned with specific biological relevance to brain tissue, rather focusing on gels that vary greatly in mesh size, the potential to alter cell phenotype, and the potential to interact with one or more of the many suspected Aβ cell surface receptors [30]. We excluded laminin and laminin-containing materials (such as Matrigel®) from these studies because of laminin’s high affinity for Aβ and its potent inhibition of fibril formation [31] Indeed, in our early experiments, we noted that Aβ did not aggregate in gels containing laminin (data not shown).

**Figure 1.**
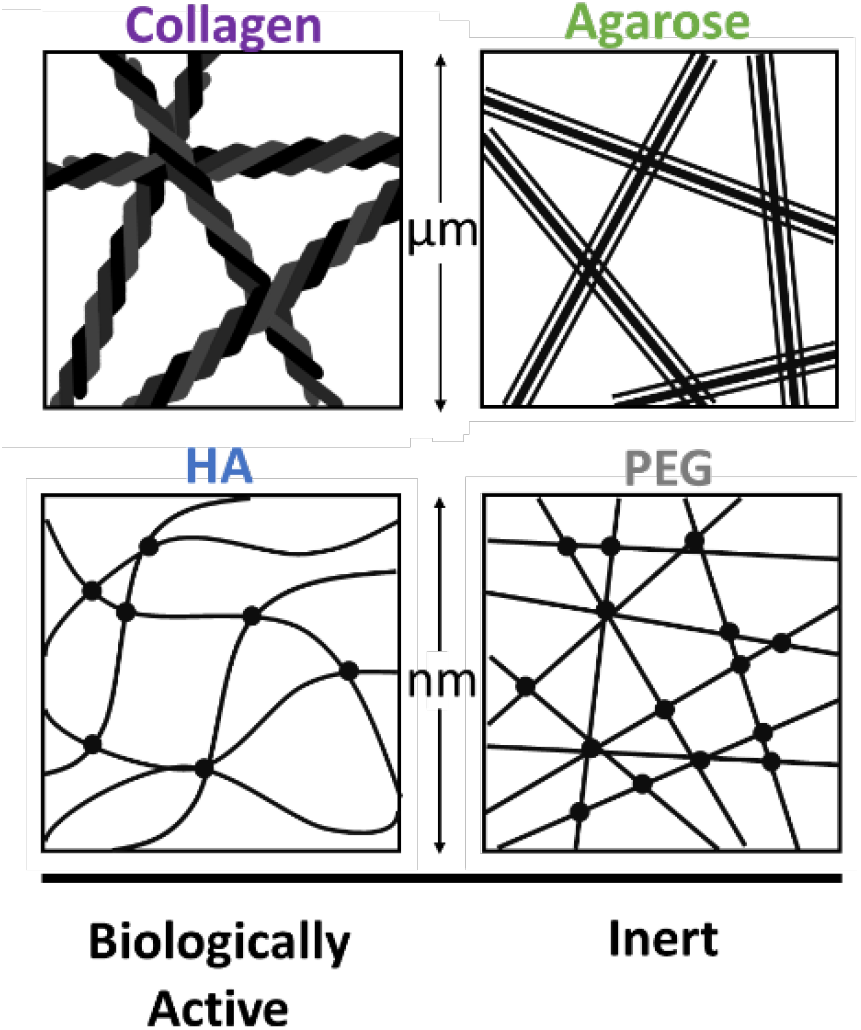
Properties of four 3D hydrogels. Four biomaterials were used as hydrogels to encapsulate PC12 cells and Aβ based on their biophysical properties. Collagen is biologically active with a mesh size ~ 10 μm. Agarose is an inert polysaccharide with a mesh size ~ 800 nm. HA is a biologically active glycosaminoglycan modified with maleimide groups and crosslinked with PEG dithiol with a mesh size ~ 200 nm. PEG is an inert polymer crosslinking a 4-arm PEG maleimide with a PEG dithiol with a mesh size ~ 10 nm.

Collagen is the most abundant ECM molecule making up 30% of total mammalian protein mass [32–33]. Type I collagen is the primary protein in the interstitial ECM and is commonly applied to *in vitro* models of cancer invasion [34–35]. Many cell types have type I collagen binding motifs that are important for adhesion, motility, and signaling [29, 36–37]. The mesh size of type I collagen hydrogels is on the order of ~10 μm [38].

Agarose is an inert polysaccharide that forms hydrogels with mesh size and stiffness that are controlled by agarose concentration and setting temperature [39]. Agarose hydrogel mesh size can range from 200 nm to 800 nm [39–40]. Agarose hydrogels have been utilized to study the diffusion of molecules through porous media [40–41] and investigate the effect of material stiffness on cell morphology [42]. In particular, pre-aggregated Aβ40 has been applied to 3D agarose culture; however, the aggregate structure was not investigated [43].

HA is a biologically-active glycosaminoglycan found in the ECM of soft connective tissues, especially the central nervous system (CNS) which is devoid of most proteinaceous ECM molecules [44–45]. Considering HA is a natural ECM molecule, it is inherently biocompatible and therefore is commonly selected for applications in regenerative medicine and drug delivery [46–48]. HA plays an important role in development and is therefore particularly relevant to *in vitro* cultures of stem cells and cancer cells [49–54]. To form stable hydrogels, HA can be modified with reactive functional groups and crosslinked to yield gels with a wide variety of properties [55–57]. HA mesh size is dependent on the molecular weight of the HA, the degree of modification of functional groups, and the crosslinking chemistry, and is typically between 100 – 600 nm [53, 58–59].

PEG is an inert synthetic polymer that can be modified with reactive functional groups and crosslinked into a hydrogel scaffold [60–61]. The particular crosslink chemistry can be selected to adjust gelation time, and the PEG molecular weight, and concentration influence gel stiffness and mesh size, which is typically 10 – 20 nm [62–63].

The work described herein examines Aβ aggregation and cytotoxicity in four hydrogels that are commonly selected for applications that involve encapsulated cells (Figure 1). We were particularly interested in collagen, agarose, HA and PEG gels because they have mesh sizes varying from ~10’s of nm to ~10’s of µm. These mesh sizes were hypothesized to impart confined microenvironments on Aβ that are relevant to the sizes of Aβ structures, from monomers/oligomers to fibrils. We were also interested in these hydrogels given their range of physiochemical properties and potential to interact with cells.

## Results and Discussion

In our earlier work, we observed that Aβ aggregation kinetics varied between the contexts of a solution and a 3D collagen hydrogel and that the variations in Aβ aggregation were associated with differences in cytotoxicity between those two contexts. We suggested that the altered Aβ aggregation in the collagen gel was due to confinement within the gel structure, which imparts a shift in the equilibrium Aβ species quickly to larger aggregates vs. the prolonged presence of oligomers in the solution of a 2D culture. Herein, we further explore this Aβ confinement effect in four hydrogel types that vary in mesh size with size scales relevant to Aβ structures, from monomers/oligomers to fibrils. Because cell-collagen-Aβ interactions may be related to the observed attenuated cytotoxicity, we were also interested in these hydrogels given their range of physiochemical properties and potential to interact with cells.

## Results

### ThT fluorescence as a measure of Aβ aggregation kinetics

To examine the impact of different 3D environments of Aβ aggregation, we used the ThT assay to identify the presence of β-sheet Aβ aggregates in solution compared to in collagen, agarose, HA, and PEG hydrogels.

Representative curves of ThT fluorescence vs. time are shown in Figure 2 for Aβ aggregation in solution and hydrogels. In solution, fibrillar Aβ aggregation (signified by ThT fluorescence) had a lag phase during the first ~20 hrs, followed by rapid aggregation. In all hydrogels, however, fibrillar aggregation did not exhibit a lag phase, and instead, fluorescence steadily increased from the initiation of the experiment (Figure 2). Depending upon the supplier and the particular lot of Aβ tested, lag time as well as the maximum fluorescence intensity varied, but all shared the same qualitative features of fibril Aβ aggregation in solution vs. the hydrogels: fibril aggregation was accelerated in the hydrogels compared to in solution. Fibrillar Aβ aggregation appeared to proceed most rapidly in the gel with the smallest mesh size -- the PEG hydrogel showed the fastest initial onset of ThT fluorescence.

**Figure 2.**
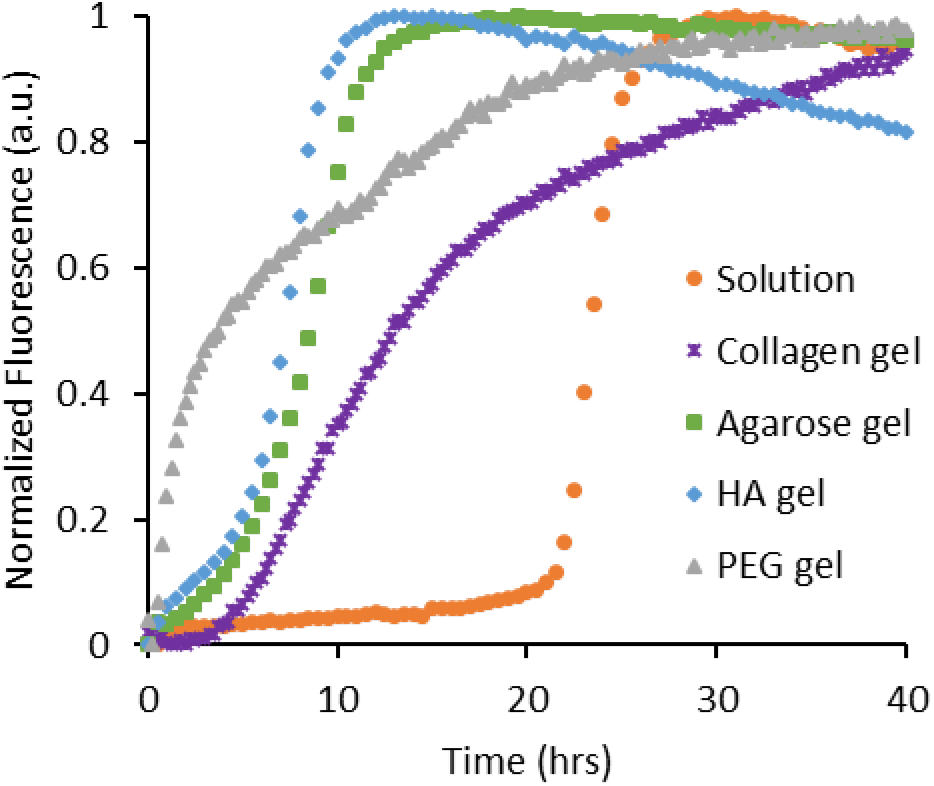
ThT Aβ aggregation kinetics in solution and four hydrogels. ThT binding to stacked β-sheet amyloids triggers fluorescence and therefore tracks kinetics of β-sheet filament aggregation in solution (orange, •), collagen hydrogel (purple, *), agarose hydrogel (green, ▪), HA hydrogel (blue, ♦), and PEG hydrogel (grey, ▴).

### Aβ aggregate diffusivities by FCS

Whereas ThT experiments provide insight into Aβ aggregate structure and kinetics, this approach is limited in that it cannot indicate aggregate size. Therefore, we utilized FCS to infer relative Aβ aggregate size from the diffusivity of fluorescently-labeled Aβ species. Diffusivity scales inversely to the radius of a spherical particle. Therefore small diffusivity values correspond to large particles. While Aβ aggregates are not spherical, this general idea that diffusivity scales inversely with particle size still applies.

Non-aggregating Scr Aβ was used as a control of monomer diffusivity. The diffusivity of these Scr Aβ monomers in solution was determined to be 175 μm^2^/s, whereas the diffusivity in the hydrogels were 129 μm^2^/s (collagen), 145 μm^2^/s (agarose), 65 μm^2^/s (HA), and 59.5 μm^2^/s (PEG) (Figure 3). As points for comparison, the diffusivity of Aβ monomer is 180 μm^2^/s in solution and 62.3 μm^2^/s in brain tissue [64]. In solution, the diffusivity of the average Aβ aggregate population (determined using the 2-component model) is ~6x slower than the monomer for up to 6 hrs (Figure 3a). In collagen, the diffusivity of the average Aβ aggregate population is ~850x slower than the monomer for up to 4 hrs (Figure 3b). In agarose, the diffusivity of the average Aβ aggregate population is ~3,600x slower than the monomer for up to 4 hrs (Figure 3c). In the small mesh size hydrogels, HA and PEG, the diffusivity of the average Aβ aggregate population are ~130x slower than the monomer with little variation (one order of magnitude or less) for up to 4 hrs (Figure 3d & e).

**Figure 3.**
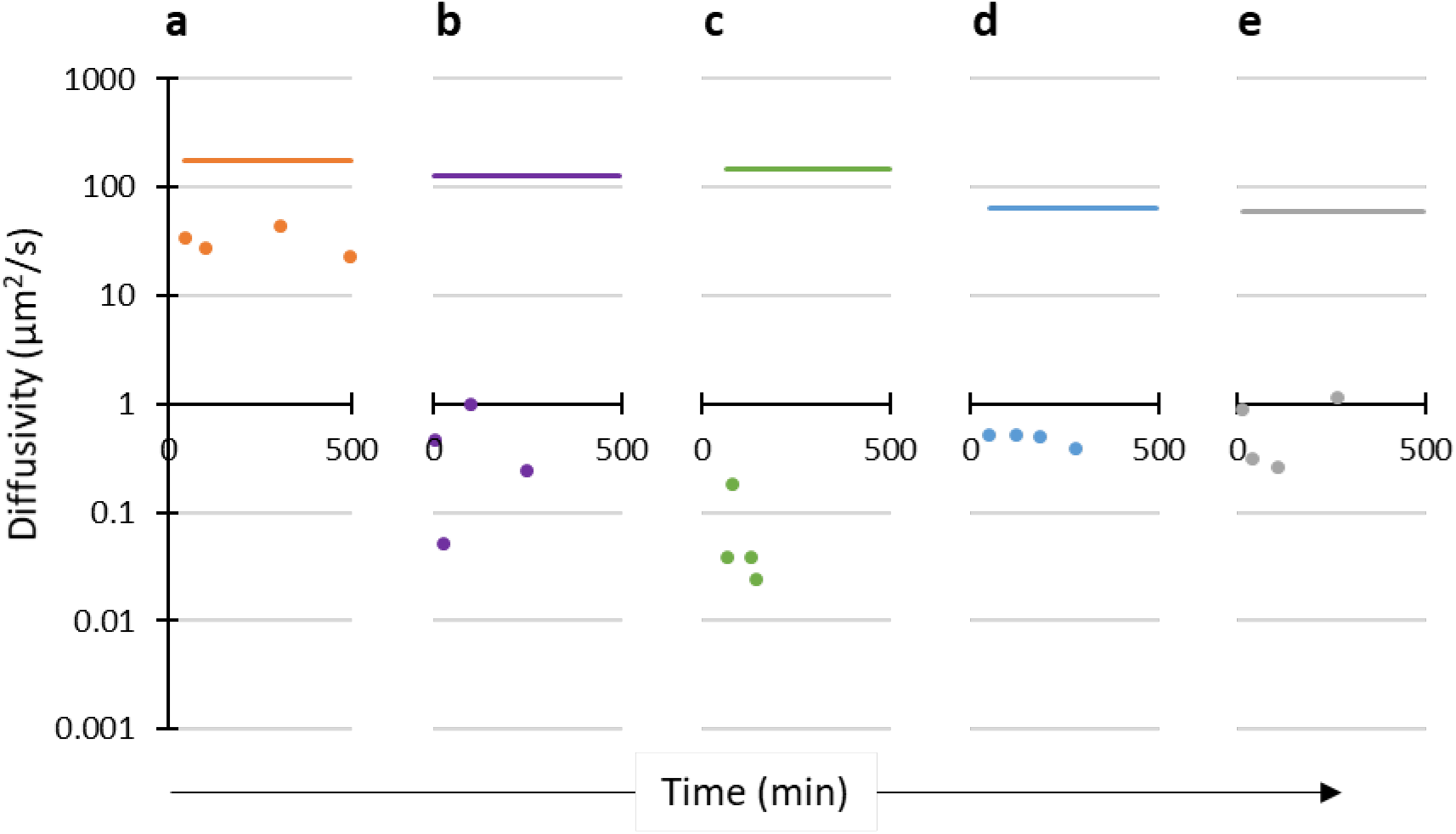
FCS G(τ) fit using a triplet 2-component model. G(τ) of 20 μM Aβ with 250 nM HiLyte Aβ where Species 1 was held constant at the calculated Scr Aβ diffusivity (in each condition) assumed to be monomer (solid line). Species 2 was solved for and represents the average aggregate species diffusivity population (•) in solution (**a**, orange), collagen hydrogel (**b**, purple), agarose hydrogel (**c**, green), HA hydrogel (**d**, blue), and PEG hydrogel (**e**, grey). The 2-sample KS test found a significant difference between all comparisons except collagen/agarose, collagen/PEG, and agarose/PEG (see Supplemental Table 1 and Supplemental Figure 1 & 2).

The correlation functions were also determined using the MEMFCS program. A distribution of multiple diffusivity populations of Aβ aggregates and their relative fractions were modeled. In solution, Aβ diffusivity values have a single broad distribution with a peak diffusivity of 85 μm^2^/s (Figure 4a). The peak diffusivity of Aβ in solution is ~2x slower than the Scr Aβ diffusivity, suggesting an Aβ population predominately composed of dimers. In all hydrogel types, Aβ has a peak diffusivity similar to the diffusivity of Scr monomer. However, in contrast to the solution samples that only have one diffusivity peak, the diffusivity values in all hydrogel types show a small secondary diffusivity peak as early as 5 mins after addition of Aβ to the hydrogel and persists throughout the measurement period (up to 4 hrs) with diffusivity values in the range of 0.17 μm^2^/s to 9 μm^2^/s, or between 360x to 50x slower than Scr Aβ (Figures 4b-e). The width of the diffusivity peaks appears to be most narrow for Aβ samples measured in the HA hydrogel.

**Figure 4.**
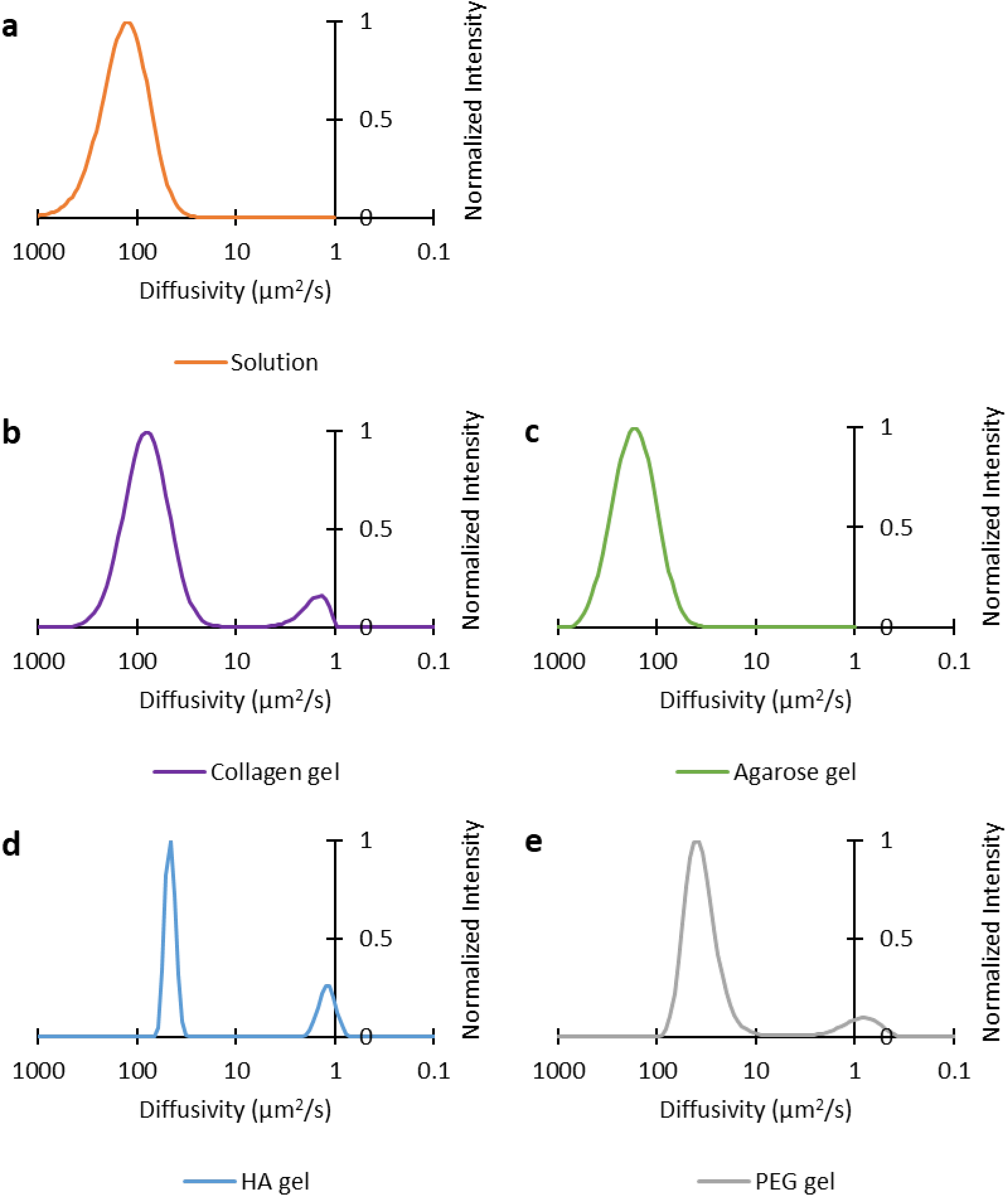
Aβ aggregate distribution using the MEMFCS program. FCS G(τ) fit of 20 μM Aβ with 250 nM HiLyte Aβ using the MEMFCS program gifted by Dr. S. Maiti [65]. In **a**, solution timepoints were collected over 8 hrs (orange; n=6). In **b**, collagen hydrogel timepoints were collected over 4 hrs (purple; n=6). In **c**, agarose hydrogel timepoints were collected over 4 hrs (green; n=5). In **d**, HA hydrogel timepoints were collected over 4 hrs (blue; n=6). In **e**, PEG hydrogel timepoints were collected over 4 hrs (grey; n=6). The 2-sample KS test found a significant difference between all comparisons except solution/agarose, collagen/PEG, and HA/PEG (see Supplemental Table 1 and Supplemental Figure 1 & 2).

Both analysis methods of the FCS data indicate that Aβ aggregates differently in 3D gels compared to in solution. Based on these data, a rough estimate of aggregate species size in hydrogels is ~25x to 200x larger than the Aβ species detected in solution.

### Toxicity of Aβ in 2D and 3D cultures

Biophysical analysis using ThT and FCS depict matching trends for Aβ aggregation in the hydrogels as compared to in solution. However, the variations in aggregation between hydrogel types may favor different size ranges of aggregate species that have varying degrees of toxicity. Therefore, we examined the viability of PC12 cells when treated with Aβ in 2D and 3D collagen, agarose, HA and PEG hydrogels over a 72 hr period (Figure 5).

**Figure 5.**
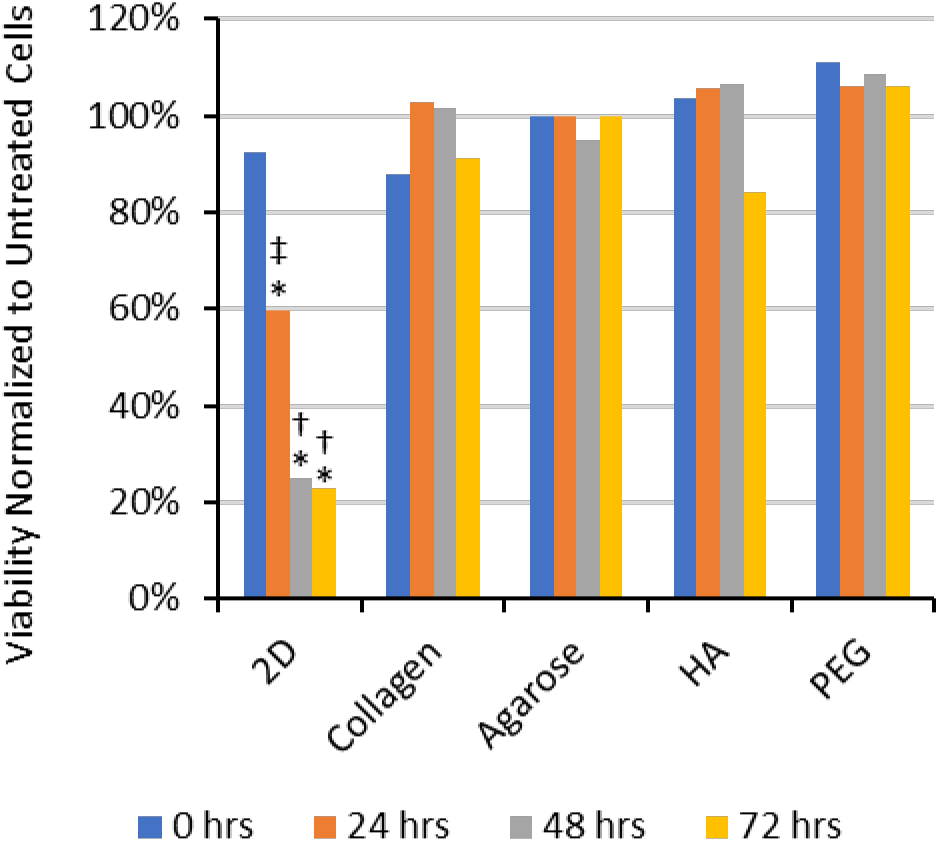
Viability of Aβ treated cells normalized by the untreated condition. Viability of PC12 cells treated with 20 μM pretreated Aβ, normalized by respective untreated conditions. Cells were cultured on 2D collagen or encapsulated within 3D collagen hydrogel, 3D agarose hydrogel, 3D HA hydrogel, or 3D PEG hydrogel. Viability was tested using a Live/Dead assay over the 72 hr period. Significant differences were seen in 2D culture in the presence of Aβ compared to no Aβ at 24, 48, and 72 hrs signified by (*). Significant differences were seen between all hydrogels treated with Aβ compared to the respective time points in 2D (48 and 72 hrs) signified by (†). A significant difference was also seen between 3D HA and 2D culture at 24 hrs signified by (‡). Statistics used n = 4. P values at significantly different times in 2D culture: 24 hrs (0.004), 48 hrs (<0.001), and 72 hrs (<0.001); 3D cultures: 48 hrs (<0.001) and 72 hrs (<0.001); and 3D HA: 24 hrs (0.045).

We acknowledge that the percent viability decreases for all samples over time, but it is important to note that the medium was not exchanged in order to better retain the evolving populations of Aβ species that were measured in the ThT and FCS experiments. Over 72 hrs, it is likely that cell waste accumulates and nutrients are depleted, thus explaining the decrease in cell viability in all conditions.

We report the viability data in two ways. In this section of the text, we report the cell viability percentages for each condition. Then to provide an alternative perspective for interpreting the results, in Figure 5, we show the same data when normalized by the respective untreated condition (Figure 5). The percent of viable cells cultured in 2D with Aβ decreased greatly by 24 hrs (49% viability; p-value 0.004), and then at 48 hrs and 72 hrs, the cell viability was further reduced to 16% (p-value <0.001) and 12% (p-value <0.001), respectively [14]. The type of 3D hydrogel did not affect cell viability, yet differences between the viability of Aβ-treated cells in 3D hydrogels vs. 2D culture at 48 hrs and 72 hrs are significant (p-value <0.001).

Considering the normalized cell viability data, at 24 hrs, cells cultured in 3D HA hydrogels treated with Aβ had higher normalized viability (105%) than in 2D (60%), and this difference is significant (p-value 0.045, Figure 5). Representative fluorescence microscopy images of LIVE/DEAD-stained cells at 0 hrs, 24 hrs, 48 hrs, and 72 hrs in 2D and 3D cultures are shown in Figure 6. In the presence of Aβ, the extent of cell death (red staining) at 48 hrs and 72 hrs in 2D culture is striking, while no notable increase in cell death is observed in the Aβ-treated 3D cultures (Figure 6b – e).

**Figure 6.**
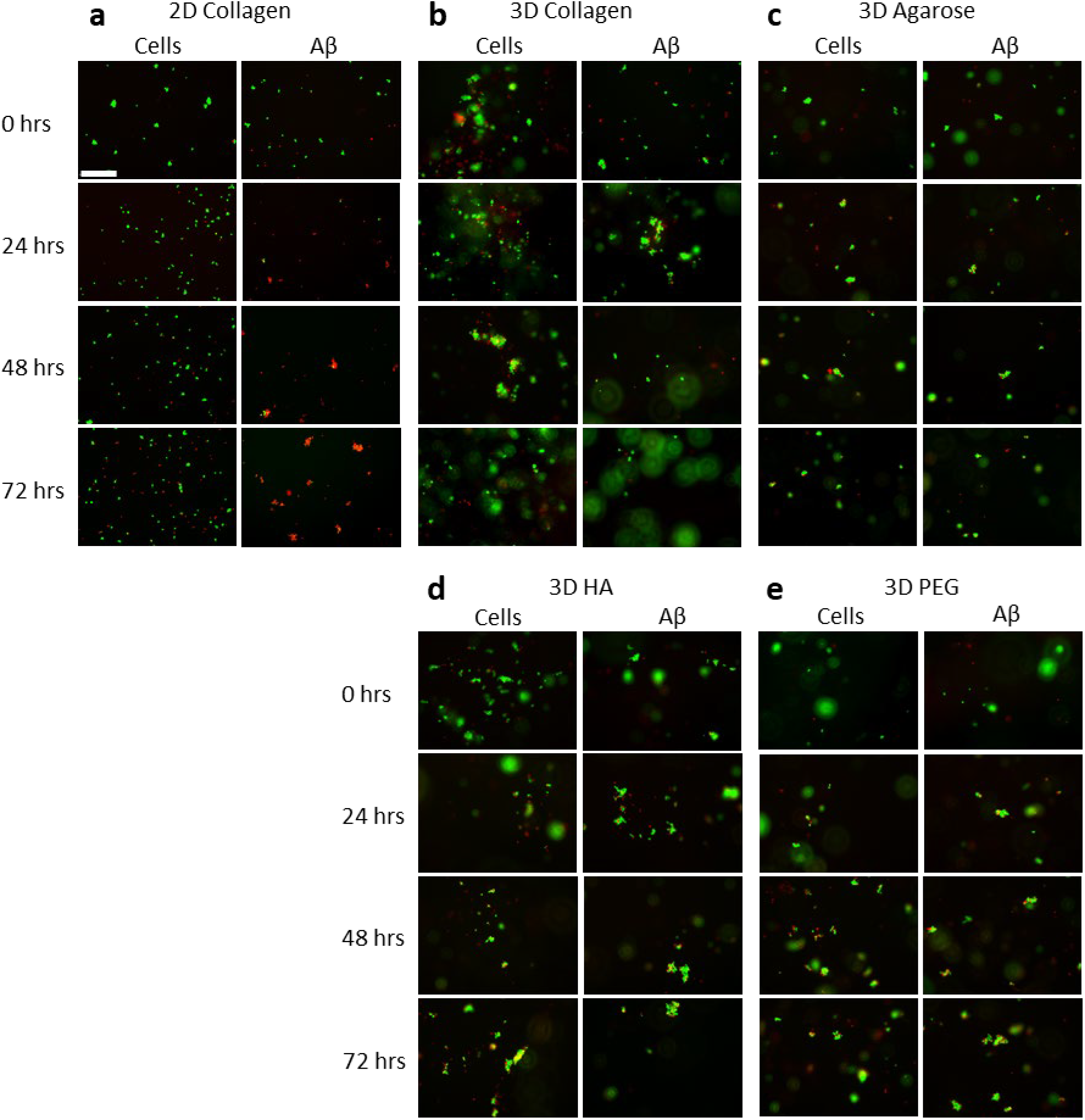
Micrographs of PC12 cell viability in 2D and 3D culture. Cells were cultured on 2D collagen (**a**), in 3D collagen (**b**), 3D agarose (**c**), 3D HA (**d**) and 3D PEG (**e**). Control conditions were cells cultured without Aβ (column labeled “Cells”). Experimental conditions were cells cultured with 20 μM Aβ (column labeled “Aβ”). Live cells fluoresced green from Calcein-AM, and dead cells fluoresced red from EthD. Scale bar is 200 μm; all micrographs are the same magnification.

## Discussion

In a broad range of contexts, the epigenetics and morphology of cells *in vivo* can be well-approximated in 3D hydrogel cultures. Though simple hydrogels cannot recapitulate the entire complexity of *in vivo* tissues, they do provide strikingly similar results compared to 2D cell culture [26–27, 29]. Depending on the application, the optimal physiochemical properties of the hydrogel model will vary. Type I collagen hydrogels have been applied to recapitulate *in vivo*-like cancer cell behaviors including migration and invasion [35, 66–67]. Agarose hydrogels are capable of allowing hepatocytes, fibroblasts, and other cell types to elaborate the distinct cellular zones that exist within respective tissues [68–70]. The bioactivity of HA hydrogels has been utilized in stem cell differentiation and patient cancer cell expansion for personalized medicine [71–72]. In addition, drug delivery applications have utilized PEG hydrogels due to their biocompatibility and tunable degradation properties [73]. Features of hydrogels that may be important in applications include mesh size, chemical composition, stiffness, and the presence of ligands or functional groups that interact with the cell surfaces.

Hydrogels impart confinement by encapsulating proteins in a macromolecular network. The network serves to exclude solvent from proteins, which minimizes the ability of proteins to undergo changes in conformation and increases the local protein concentration; the net result is that confinement promotes proteins to undertake compact structures and favors protein-protein interactions. The degree to which a particular hydrogel exerts confinement on an encapsulated protein is inversely proportional to the gel’s mesh size. With this in mind, we predicted altered Aβ aggregation kinetics in gels with small mesh sizes (HA, PEG) vs. those with larger mesh sizes (collagen, agarose).

The attenuation of Aβ toxicity in hydrogels could potentially be influenced by cellular changes that occur due to cell-hydrogel interactions. Thus, we acknowledge that the biological activity of the hydrogel (e.g., integrin-binding motifs in collagen and CD44- and RHAMM-binding motifs in HA) may influence Aβ toxicity. Collagen is a commonly used hydrogel, but HA hydrogels could be more relevant to studies related to the brain wherein the main ECM molecules are HA, tenascins, and lecticans [45]. Basement membrane ECM molecules such as laminins and collagens are an important component of the blood-brain barrier and are found surrounding blood vessels in the brain [74]. Thus, we investigated Aβ aggregation and cytotoxicity in both biologically-active and inert hydrogels in order to uncover a possible role of bioactivity on cell susceptibility to Aβ toxicity.

The kinetics of Aβ aggregation was measured via ThT assay, wherein ThT fluorescence indicates the presence of β-sheet structures. The presence of β-sheet aggregates was negligible (no fluorescence) at the start of each experiment. This is consistent with the Aβ pretreatment process that was used to ensure a consistent population of Aβ monomers at the start of each experiment [14]. All conditions showed the presence of aggregated Aβ (ThT fluorescence) that increased over time according to one of four trends: 1) a pronounced lag phase then rapid aggregation, 2) a two-phase increase in fluorescence depicting an overall relatively slow rate of aggregation, 3) a brief slow phase then rapid aggregation, and 4) a two-phase increase in fluorescence suggesting an overall relatively fast rate of aggregation.

For Aβ aggregation in solution, ThT fluorescence measurements consistently show a pronounced lag phase then rapid aggregation or a trend 1 curve (Figure 2). Both ThT and FCS measurements indicate that the only Aβ species present for up to 20 hrs were small, rapidly diffusing species that are devoid of β-sheet structures for at least 6 hrs (Figures 2, 3a & 4a). At these early times, the Aβ species present are likely monomers and dimers as well as a population of larger species (diffusivities of ~20 μm^2^/s, Figure 3a, 4a) do not have extended β-sheet structure (do not bind ThT, Figure 2)

For Aβ aggregation in 3D hydrogels, all four types displayed the immediate presence of extended β-sheet structures and a population of large aggregate species (Figure 2, 3b-e & 4b-e). In other words, for all hydrogels, a type 1 curve was never observed, but instead displayed trend 2, 3, or 4 with some differences in magnitude depending on Aβ lot and random variation. Figure 2 depicts the most common curves observed for each hydrogel type: collagen shows trend 2, agarose shows trend 3, HA shows trend 3, and PEG shows trend 4. These trends are consistent with the idea that Aβ is confined in a hydrogel: PEG hydrogels have the smallest mesh size (~20 nm) and show the fastest Aβ aggregation, collagen hydrogels have the largest mesh size (~10 μm) and show Aβ aggregation occurring at a slower rate than in PEG gels, yet faster than in solution.

We expected that given the differences in the Aβ aggregation rate and size distribution observed in the four hydrogel types, coupled with differences in hydrogel bioactivity, that Aβ cytotoxicity would vary with the particular hydrogel type, but all hydrogels would be associated with lower cytotoxicity than that observed in solution. To our surprise, despite quantitative differences in Aβ aggregation kinetics and aggregate size distributions, all hydrogel materials completely attenuated Aβ toxicity for up to 72 hrs in culture (Figure 5 & 6). When Aβ aggregation was reexamined in hydrogels containing cells, there was no difference in respective aggregation kinetic curve types regardless of the presence of cells (data not shown). From the results presented herein, it appears the key feature relevant to Aβ toxicity that is consistent across all hydrogel types is the rapid stabilization of large β-sheet aggregates, suggesting that attenuated Aβ cytotoxicity in hydrogels may be due to a limited presence of Aβ oligomers that are available to interact with cells.

We acknowledge that Aβ aggregation also may be influenced by properties of the hydrogels (e.g., charge, hydrophilicity) that were not evaluated herein. Also, we cannot rule out that the attenuation of Aβ toxicity in hydrogels is influenced by confinement of the cells themselves or effects from the stiffness of the cellular microenvironment. However, we hold that these possibilities are unlikely given the evidence that 3D culture allows cells to more closely mimic *in vivo* phenotypes vs. 2D culture [26–27, 29]. More importantly, we reported in 2002 that agarose itself does not confer a protective effect against the cytotoxicity of the Aβ 1-40 amino acid sequence [43]. Similar results were found with the 1-42 amino acid sequence of Aβ (unpublished data). Results presented herein may seem to contradict findings in our 2002 publication [43] that Aβ is toxic to cells in 3D agarose hydrogels. It is important to note that experiments herein utilized Aβ in the monomeric form, whereas the 2002 publication used pre-aggregated Aβ that contained a mixture of fibrils and smaller aggregated species, including the toxic 20 µm^2^/s diffusing species. In contrast, we report here that the intermediate 20 µm^2^/s diffusing species was not observed in any of the hydrogel types tested (Figures 3 & 4). Therefore, the current and 2002 reports are consistent in the idea that confinement in a hydrogel alters the kinetics of Aβ aggregation resulting in a) Aβ populations predominated by larger aggregate species (as opposed to Aβ in solution wherein oligomers are present for prolonged times), and b) attenuation of Aβ toxicity vs that observed 2D cultures.

Our findings have strong implications for *in vitro* models of disease. Aβ has been studied for decades in solution; wherein unstructured cytotoxic aggregates are clearly identifiable. Many drugs have been designed to target Aβ aggregation or interactions with cells. Yet, astoundingly few AD drugs have been approved by the FDA. We demonstrate here that Aβ cytotoxicity is completely attenuated in 3D culture models composed of commonly-used hydrogels that have a broad range of physical, chemical and biological properties.

Stated more generally, we report that protein-protein interactions are altered in confined microenvironments. We suggest that this phenomenon may also relate to protein confinement as it occurs intracellularly and *in vivo.* Therefore, any field of research investigating protein structure and function in contexts relevant to those that exist *in vivo* should consider the potential impact of protein confinement by the local microenvironment.

## Materials and Methods

### Beta-Amyloid Preparation

Human beta-amyloid (1-42) (Aβ) and scrambled Aβ (1-42) (Scr Aβ) (*AIAEGDSHVLKEGAYMEIFDVQGHVFGGKIFRVVDLGSHNVA*) was purchased from AnaSpec (Fremont, CA) and Genscript (Piscataway, NJ). HiLyte 488-labeled Aβ (1-42) (HiLyte Aβ) and FAM-labeled scrambled Aβ (1-42) (FAM Scr Aβ) were purchased from AnaSpec (Fremont, CA). All other unspecified reagents were purchased from Sigma Aldrich (St. Louis, MO) or ThermoFisher Scientific (Waltham, MA).

To break any existing β-sheet structures and monomerize the protein, lyophilized Aβ was pretreated with hexafluoro-2-propanol at a concentration of 1 mg/ml for 40 mins until Aβ was fully dissolved. Aβ aliquots were transferred into glass scintillation vials, and hexafluoro-2-propanol was evaporated under vacuum overnight. Aliquots of dried peptide film were stored at −20°C. For an experiment, an Aβ aliquot was dissolved in freshly-made and filtered 60 mM NaOH and allowed to dissolve for 2 mins at room temperature. Tissue culture grade water was then added, and the vial was sonicated for 5 mins. Next, the Aβ solution was filtered with a 0.2-μm pore, 4-mm diameter syringe filter. Sterile phosphate buffered saline (PBS) was then added to the Aβ monomer solution yielding a final concentration of 222 μM with the NaOH:water:PBS ratio of 2:7:1. The Aβ solution was used immediately after preparation. HiLyte Aβ and FAM Scr Aβ were prepared in the same NaOH:water:PBS ratio solution to a stock Aβ concentration of 10 μM.

### Hydrogel Preparation

Rat tail type I collagen hydrogels were prepared to final concentrations of 1 mg/ml. Cold 5x Dulbecco’s Modified Eagle’s Medium (DMEM) without phenol red, 7.5% sodium bicarbonate, sterile deionized water, and collagen were combined with PC12 cells to generate 3D substrates in black-walled clear-bottom well plates.

SeqPlaque Agarose with a concentration of 1% (w/v) was prepared in deionized water and sterilized. Agarose was heated to 68°C then cooled at room temperature for 5 minutes before mixing 1:1 with concentrated culture medium; yielding a solution of 1x DMEM without phenol red, 1% B27, and 0.5% agarose. The hydrogel solution was dispensed into black-walled clear-bottom well plates and placed in a culture incubator for 20 mins to allow for gelation.

HA and PEG hydrogels were each crosslinked by a maleimide-thiol Michael addition click reaction. Hyaluronic acid (HA, 242 kDa) was functionalized with maleimide (HA-Mal) following a published protocol [59]. Briefly, HA was dissolved in 0.1 M 2-(N-morpholino)ethanesulfonic acid (MES) buffer at a concentration of 5.15 mM. 1-Ethyl-3-(3-dimethylaminopropyl) carbodiimide (EDC, 15 mM) and N-hydroxysuccinimide (NHS, 15 mM) were added, and the solution was mixed for 30 mins. Next, N-(2-aminoethyl) maleimide trifluoroacetate salt (AEM, 10 mM) was added and mixed for 4 hrs covered with plastic wrap. The mixture was dialyzed against 50 mM NaCl in deionized water for 3 days, then against deionized water for 3 days. The dialyzed solution was then sterile-filtered and aliquoted aseptically into sterile 15-ml tubes, lyophilized and stored at - 20°C. The degree of substitution, the number of maleimide groups per HA chain, was determined as per [59] by ^1^H NMR to be ~40.

For HA hydrogels, HA-Mal was prepared at 1% (w/v) and mixed equal volume with PEG dithiol (10 kDa) at a molar ratio of 1:1.2 maleimide to thiol. For PEG hydrogels, 4-arm PEG maleimide (PEG-Mal, 20 kDa) was prepared at 5% (w/v) and mixed equal volume with PEG dithiol (10 kDa) at a molar ratio of 1:1 maleimide to thiol. All HA and PEG solutions were dissolved in Neurobasal medium supplemented with 1% B27. PEG solutions were filter-sterilized. In black-walled clear-bottom well plates, maleimide solutions were pipetted into the well first, then the thiol solution containing experimental additives (cells, Aβ, ThT) was pipetted into the maleimide droplet to mix. Both HA and PEG gels crosslinked within ~5 sec.

### Thioflavin T

A black-walled clear-bottom 384 well plate (Costar) was sterilized under UV light for 15 minutes in a laminar flow hood. UltraPure grade Thioflavin T (ThT) (AnaSpec, Fremont, CA) was dissolved in deionized water at a concentration of 1 mM then filter-sterilized. Wells for 2D and 3D samples were prepared as above but contained 20 µM ThT. The wells were sealed with black TopSeal-A membranes to prevent evaporation. The ThT experiment was analyzed on a Spectra Max M5 (Molecular Devices, San Jose, CA) spectrophotometer set to ex. 450 nm, em. 480 nm, at 37°C, taking measurements every 30 mins for 72 hrs and reading from the bottom of the plate. Replicates were averaged, Aβ data was corrected with ThT control data, and corrected curves were normalized. As in standard practice in Aβ aggregation studies [75–76] due to the stochastic nature of aggregation, curves representative of at least 10-20 experiments are presented here.

### Fluorescence Correlation Spectroscopy

#### Theory

Fluorescence Correlation Spectroscopy (FCS) measures the fluctuations of fluorescence in a small, optically-defined confocal volume (~10^−15^ liter). These fluctuations are typically attributed to the fluorescent particles moving in and out of the volume with a statistical average residence time, τ_D_. The residence time is proportional to the hydrodynamic radius (R_H_) of the molecule. The fluctuations of detected photons inform the autocorrelation, G(τ), function defined as

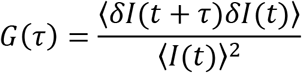

Where *δI*(*t*) = *I*(*t*) − 〈*I*(*t*)〉 is the fluorescence fluctuation determined from the measured fluorescence intensity, (*t*), at time t, and the average intensity, 〈*I*(*t*)〉, over the period of measurement. The excitation laser, which is focused, is assumed to have a 3D Gaussian profile, with a characteristic radial dimension (w_0_) and a characteristic axial dimension (z_0_). For a solution of n noninteracting, freely diffusing fluorescent species G(τ) is given by:

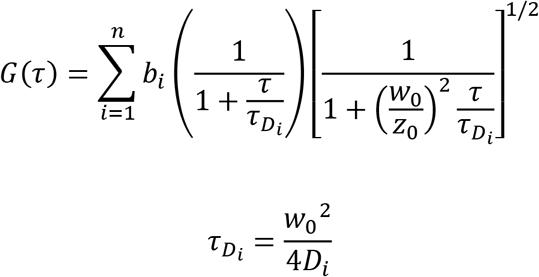

Here the D_i_ values are the n different values of diffusion constants and b_i_ are the relative fractions in brightness of these species. In practice, the radial and axial dimensions were determined using Alexa 488 dye in water where the diffusion coefficient (430 μm^2^/s) is known and was used to estimate the excitation volume for a 3D Gaussian beam [77].

#### Methods

Neurobasal medium was used in preparing solution samples and contained 20 μM Aβ and 250 nM HiLyte Aβ. Hydrogels were prepared as described with 20 μM Aβ and 250 nM HiLyte Aβ then pipetted into 0.8-mm deep hybridization chambers (PerkinElmer, Waltham, MA) on a borosilicate cover glass. Control samples were tested with 20 μM Scr Aβ and 250 nM FAM Scr Aβ.

The FCS measurements were performed using an Alba-FFS microscope-based system from ISS Inc. (Champagne, IL). The system is composed of: an Olympus IX81 inverted microscope equipped with a 60X/1.35NA oil immersion objective lens, a Prior Pro stage, three different lasers (450 nm, 488 nm, and 532 nm), two Hamamatsu Photon Multiplier tubes (PMTs) for photodetection, and two sets of computer-controlled scanned mirrors for imaging. In these measurements, only the 488-nm diode laser was used for excitation of the fluorophores Alexa 488 or fluorescently-labeled Aβ, and the emitted fluorescence was collected through confocal detection with a pinhole (< 50 mm) located in the image plane of the excited focused beam inside the sample. The emitted fluorescent beam was optically filtered further with (525/50nm) filter and then sent to a 50/50 beam splitter for detection by two PMTs positioned in a 90-degree angle configuration. The photocounts of both PMTs were continuously acquired and then computationally cross-correlated in order to eliminate the afterpulsing effect of a single PMT, which is typically noticeable at short delay times (< 10 ms).

Using Vista Vision software, two runs were carried out back to back collecting for 3 minutes each to generate the correlation function G(τ) for each sample at a time point. The two correlation functions were averaged, and the Scr Aβ correlation function was fit using the one-component model to determine the diffusivity of the monomer. Further, the measured time-correlation functions for Aβ were fit using the 2-component model where the size of species 1 was held constant at monomer diffusivity in order to derive the average aggregate diffusivity population of the second species. Additional refinement for fitting the correlation functions were also performed with the Maximum Entropy Method FCS (MEMFCS) thanks to a code gifted by Dr. S. Maiti (Tata Institute of Fundamental Research), allowing us to obtain the heterogeneous distribution of aggregate diffusivities at each time point [78].

Small molecules have a short delay time because they diffuse quickly through the volume, whereas large molecules have a long delay time because of their relatively slow diffusion through the volume. The 2-component model is intended to model two distinct molecular species in solution. For our samples, we held the monomer diffusivity constant as species 1 where the average diffusivity of aggregated species was identified by solving for species 2.

Fluorophore labeling of Aβ monomers inhibits aggregation due to the bulky groups sterically preventing proper monomer to monomer stacking [79]. Therefore, we used a ratio of 1:80 HiLyte 488-labeled Aβ to unlabeled Aβ, and FAM-labeled Scr Aβ to unlabeled Scr Aβ, to allow unhindered β-sheet stacking. Nanomolar fluorophore concentrations are also preferable in FCS in order for the detectors to monitor few individual fluorescent molecules in the confocal volume, enhancing hence the signal-to-noise of the fluctuations.

### Cell Culture

PC12 cells (ATCC, Manassas, VA) (CRL-1721TM) were cultured in collagen-coated flasks. Growth medium consisted of DMEM/F12 with L-glutamine and without phenol red, supplemented with 10% inactivated horse serum, 5% fetal bovine serum, and 20 μg/ml gentamicin. Experimental medium consisted of Neurobasal medium without phenol red, supplemented with 1% B27 and 20 μg/ml gentamicin. Phenol red and serum were avoided in the experiments because they are inhibitors of Aβ aggregation [80–81].

### Live/Dead Assay

PC12 cells were collected by trypsin treatment, and viability was determined by trypan blue staining. To remove serum, cells were resuspended in experimental medium, pelleted then resuspended again in experimental media. In a black-walled clear-bottom tissue culture treated 96-well plate, wells for 2D culture were collagen-coated, and then PC12 cells were seeded at 15 × 10^3^ cell/cm^2^. For the 3D hydrogels, PC12 cells were mixed in collagen and agarose gel solution at a concentration of 500 cell/μl; the solution was then pipetted (30 μl) into the well plate and allowed to solidify. For HA and PEG hydrogels, PC12 cells were mixed in PEG dithiol solutions at a concentration of 1000 cell/μl. HA-Mal and PEG-Mal solutions were pipetted (15 μl) into the well first; then the PEG dithiol solution (containing cells) was pipetted (15 μl) into the maleimide solution to mix. The final HA and PEG hydrogels had a PC12 cell concentration of 500 cell/μl. All wells were incubated in 200 µl warmed medium.

To determine cell viability, the Live/Dead mammalian cell kit (Invitrogen, Carlsbad, CA) was applied at a concentration of 4 μM Calcein AM (green-fluorescing live cell reporter) and 9 μM Ethidium homodimer-1 (EthD) (red-fluorescing dead cell reporter) and incubated at 37°C for 30 minutes. Images were captured on an IX81 Olympus inverted fluorescent microscope. A minimum of 100 cells were counted per well (two images per well), and three wells per condition were tested. The data is presented as percent viability, averaged between the three replicate experiments.

### Statistical analysis

Data were analyzed for statistical significance with Prism v8 software (GraphPad). The raw FCS experimental G(τ) curves, the 2-component model calculated G(τ) curves, and the MEMFCS calculated G(τ) curves were analyzed for significance using the 2-sample Kolmogorov-Smirnov test with 95% confidence. To correct for multiple comparisons, we used the two-stage step-up method of Benjamini, Krieger and Yekutieli with the false discovery rate (FDR) set to 5%. Cell viability data were analyzed with a general ANOVA with a post Tukey pairwise test which determined significant deviation from the population mean with a p-value <0.05 with 95% confidence.

## Abbreviations

AD: Alzheimer’s disease
Aβ: amyloid-β
DMEM: Dulbecco’s Modified Eagle’s Medium
ECM: extracellular matrix
FDR: false discovery rate
FAM Scr Aβ: FAM-labeled scrambled Aβ (1-42)
FCS: fluorescence correlation spectroscopy
HiLyte Aβ: HiLyte 488-labeled Aβ (1-42)
HA: hyaluronic acid
MEMFCS: Maximum Entropy Method FCS
NHS: N-hydroxysuccinimide
PBS: phosphate buffered saline
PEG: polyethylene glycol
ThT: thioflavin T

## Author Contributions

LWS, TAG and JBL developed the concept and designed experiments. LWS performed all experiments and data analysis. GLS contributed towards statistical analysis of the FCS data. HB contributed towards FCS data acquisition and provided FCS instrumentation. LWS wrote the manuscript text with editing comments by TAG and JBL.

## Acknowledgments

The authors would like to thank Tagide deCarvalho for her assistance with TEM imaging and Dr. S. Maiti (Tata Institute of Fundamental Research) for sharing the Maximum Entropy Method program for FCS. This work was supported by funding from NSF (EAGER CBET-1447057) and NIH (R01GM117159). NSF provided support for TAG to contribute to this project through their Independent Research and Development program. Any opinion, findings, and conclusions or recommendations expressed in this material are those of the author(s) and do not necessarily reflect the views of the National Science Foundation.

## References

1. Khan, F.; Tanaka, M., Designing Smart Biomaterials for Tissue Engineering. Int J Mol Sci 2017, 19

2. Walsh, D. M.; Selkoe, D. J., A beta oligomers - a decade of discovery. J Neurochem 2007, 101 (5), 1172–84.

3. Lee, S.; Fernandez, E. J.; Good, T. A., Role of aggregation conditions in structure, stability, and toxicity of intermediates in the Abeta fibril formation pathway. Protein Sci 2007, 16 (4), 723–32.

4. Glabe, C. G., Structural classification of toxic amyloid oligomers. J Biol Chem 2008, 283 (44), 29639–43.

5. Ahmed, M.; Davis, J.; Aucoin, D.; Sato, T.; Ahuja, S.; Aimoto, S.; Elliott, J. I.; Van Nostrand, W. E.; Smith, S. O., Structural conversion of neurotoxic amyloid-beta(1-42) oligomers to fibrils. Nat Struct Mol Biol 2010, 17 (5), 561–7.

6. Cizas, P.; Budvytyte, R.; Morkuniene, R.; Moldovan, R.; Broccio, M.; Losche, M.; Niaura, G.; Valincius, G.; Borutaite, V., Size-dependent neurotoxicity of beta-amyloid oligomers. Arch Biochem Biophys 2010, 496 (2), 84–92.

7. Dubnovitsky, A.; Sandberg, A.; Rahman, M. M.; Benilova, I.; Lendel, C.; Hard, T., Amyloid-beta protofibrils: size, morphology and synaptotoxicity of an engineered mimic. PLoS One 2013, 8 (7), e66101.

8. Tanzi, R. E.; Bertram, L., Twenty years of the Alzheimer’s disease amyloid hypothesis: a genetic perspective. Cell 2005, 120 (4), 545–55.

9. Cummings, J.; Lee, G.; Mortsdorf, T.; Ritter, A.; Zhong, K., Alzheimer’s disease drug development pipeline: 2017. Alzheimers Dement (N Y) 2017, 3 (3), 367–384.

10. Cummings, J.; Morstorf, T.; Lee, G., Alzheimer’s drug-development pipeline: 2016. Alzheimers Dement (N Y) 2016, 2 (4), 222–232.

11. Schneider, L. S.; Mangialasche, F.; Andreasen, N.; Feldman, H.; Giacobini, E.; Jones, R.; Mantua, V.; Mecocci, P.; Pani, L.; Winblad, B.; Kivipelto, M., Clinical trials and late-stage drug development for Alzheimer’s disease: an appraisal from 1984 to 2014. J Intern Med 2014, 275 (3), 251–83.

12. Banik, A.; Brown, R. E.; Bamburg, J.; Lahiri, D. K.; Khurana, D.; Friedland, R. P.; Chen, W.; Ding, Y.; Mudher, A.; Padjen, A. L.; Mukaetova-Ladinska, E.; Ihara, M.; Srivastava, S.; Padma Srivastava, M. V.; Masters, C. L.; Kalaria, R. N.; Anand, A., Translation of Pre-Clinical Studies into Successful Clinical Trials for Alzheimer’s Disease: What are the Roadblocks and How Can They Be Overcome? J Alzheimers Dis 2015, 47 (4), 815–43.

13. Servick, K., Another major drug candidate targeting the brain plaques of Alzheimer’s disease has failed. What’s left? Science 2019, 363 (6433).

14. Simpson, L. W.; Hacene, B.; Good, T. A.; Leach, J. B., Collagen hydrogel confinement of Amyloid-β (Aβ) accelerates aggregation and reduces cytotoxic effects. Submitted to Journal on Date 2019.

15. Choi, S. H.; Kim, Y. H.; Hebisch, M.; Sliwinski, C.; Lee, S.; D’Avanzo, C.; Chen, H.; Hooli, B.; Asselin, C.; Muffat, J.; Klee, J. B.; Zhang, C.; Wainger, B. J.; Peitz, M.; Kovacs, D. M.; Woolf, C. J.; Wagner, S. L.; Tanzi, R. E.; Kim, D. Y., A three-dimensional human neural cell culture model of Alzheimer’s disease. Nature 2014, 515 (7526), 274–8.

16. Park, J.; Wetzel, I.; Marriott, I.; Dreau, D.; D’Avanzo, C.; Kim, D. Y.; Tanzi, R. E.; Cho, H., A 3D human triculture system modeling neurodegeneration and neuroinflammation in Alzheimer’s disease. Nat Neurosci 2018, 21 (7), 941–951.

17. Jorfi, M.; D’Avanzo, C.; Tanzi, R. E.; Kim, D. Y.; Irimia, D., Human Neurospheroid Arrays for In Vitro Studies of Alzheimer’s Disease. Sci Rep 2018, 8 (1), 2450.

18. Ghourichaee, S. S.; Powell, E. M.; Leach, J. B., Enhancement of human neural stem cell self-renewal in 3D hypoxic culture. Biotechnol Bioeng 2017, 114 (5), 1096–1106.

19. Yagi, T.; Ito, D.; Okada, Y.; Akamatsu, W.; Nihei, Y.; Yoshizaki, T.; Yamanaka, S.; Okano, H.; Suzuki, N., Modeling familial Alzheimer’s disease with induced pluripotent stem cells. Hum Mol Genet 2011, 20 (23), 4530–9.

20. Luo, Y.; Lou, C.; Zhang, S.; Zhu, Z.; Xing, Q.; Wang, P.; Liu, T.; Liu, H.; Li, C.; Shi, W.; Du, Z.; Gao, Y., Three-dimensional hydrogel culture conditions promote the differentiation of human induced pluripotent stem cells into hepatocytes. Cytotherapy 2018, 20 (1), 95–107.

21. Wang, B.; Jakus, A. E.; Baptista, P. M.; Soker, S.; Soto-Gutierrez, A.; Abecassis, M. M.; Shah, R. N.; Wertheim, J. A., Functional Maturation of Induced Pluripotent Stem Cell Hepatocytes in Extracellular Matrix-A Comparative Analysis of Bioartificial Liver Microenvironments. Stem Cells Transl Med 2016, 5 (9), 1257–67.

22. Ghourichaee, S. S.; Leach, J. B., The effect of hypoxia and laminin-rich substrates on the proliferative behavior of human neural stem cells. J Mater Chem B 2016, 4 (20), 3509–3514.

23. Postovit, L. M.; Seftor, E. A.; Seftor, R. E.; Hendrix, M. J., A three-dimensional model to study the epigenetic effects induced by the microenvironment of human embryonic stem cells. Stem Cells 2006, 24 (3), 501–5.

24. Engler, A. J.; Sen, S.; Sweeney, H. L.; Discher, D. E., Matrix elasticity directs stem cell lineage specification. Cell 2006, 126 (4), 677–89.

25. Cavo, M.; Caria, M.; Pulsoni, I.; Beltrame, F.; Fato, M.; Scaglione, S., A new cell-laden 3D Alginate-Matrigel hydrogel resembles human breast cancer cell malignant morphology, spread and invasion capability observed “in vivo”. Sci Rep 2018, 8 (1), 5333.

26. Mirbagheri, M.; Adibnia, V.; Hughes, B. R.; Waldman, S. D.; Banquy, X.; Hwang, D. K., Advanced cell culture platforms: a growing quest for emulating natural tissues. Mater Horiz 2019, 6 (1), 45–71.

27. Balasubramanian, S.; Packard, J. A.; Leach, J. B.; Powell, E. M., Three-Dimensional Environment Sustains Morphological Heterogeneity and Promotes Phenotypic Progression During Astrocyte Development. Tissue Eng Part A 2016, 22 (11-12), 885–98.

28. Ribeiro, A.; Balasubramanian, S.; Hughes, D.; Vargo, S.; Powell, E. M.; Leach, J. B., beta1-Integrin cytoskeletal signaling regulates sensory neuron response to matrix dimensionality. Neuroscience 2013, 248, 67–78.

29. Ribeiro, A.; Vargo, S.; Powell, E. M.; Leach, J. B., Substrate three-dimensionality induces elemental morphological transformation of sensory neurons on a physiologic timescale. Tissue Eng Part A 2012, 18 (1-2), 93–102.

30. Jarosz-Griffiths, H. H.; Noble, E.; Rushworth, J. V.; Hooper, N. M., Amyloid-beta Receptors: The Good, the Bad, and the Prion Protein. J Biol Chem 2016, 291 (7), 3174–83.

31. Castillo, G. M.; Lukito, W.; Peskind, E.; Raskind, M.; Kirschner, D. A.; Yee, A. G.; Snow, A. D., Laminin inhibition of β-amyloid protein (Aβ) fibrillogenesis and identification of an Aβ binding site localized to the globular domain repeats on the laminin a chain. J. Neurosci. Res. 2000, 62 (3), 451–462.

32. Myllyharju, J.; Kivirikko, K. I., Collagens, modifying enzymes and their mutations in humans, flies and worms. Trends Genet 2004, 20 (1), 33–43.

33. Shoulders, M. D.; Raines, R. T., Collagen structure and stability. Annu Rev Biochem 2009, 78, 929–58.

34. Boot-Handford, R. P.; Tuckwell, D. S., Fibrillar collagen: the key to vertebrate evolution? A tale of molecular incest. Bioessays 2003, 25 (2), 142–51.

35. Rianna, C.; Kumar, P.; Radmacher, M., The role of the microenvironment in the biophysics of cancer. Semin Cell Dev Biol 2017.

36. Parkhurst, M. R.; Saltzman, W. M., Quantification of human neutrophil motility in three-dimensional collagen gels. Effect of collagen concentration. Biophys J 1992, 61 (2), 306–315.

37. Baker, B. M.; Chen, C. S., Deconstructing the third dimension: how 3D culture microenvironments alter cellular cues. J Cell Sci 2012, 125 (Pt 13), 3015–24.

38. Banerjee, P.; Lenz, D.; Robinson, J. P.; Rickus, J. L.; Bhunia, A. K., A novel and simple cell-based detection system with a collagen-encapsulated B-lymphocyte cell line as a biosensor for rapid detection of pathogens and toxins. Lab Invest 2008, 88 (2), 196–206.

39. Narayanan, J.; Xiong, J.-Y.; Liu, X.-Y., Determination of agarose gel pore size: Absorbance measurements vis a vis other techniques. J Phys Conf Ser. 2006, 28, 83–86.

40. Pluen, A.; Netti, P. A.; Jain, R. K.; Berk, D. A., Diffusion of Macromolecules in Agarose Gels: Comparison of Linear and Globular Configurations. Biophys J 1999, 77 (1), 542–552.

41. Johnson, E. M.; Berk, D. A.; Jain, R. K.; Deen, W. M., Diffusion and partitioning of proteins in charged agarose gels. Biophys J 1995, 68 (4), 1561–1568.

42. Balgude, A. P.; Yu, X.; Szymanski, A.; Bellamkonda, R. V., Agarose gel stiffness determins rate of DRG neurite extension in 3D cultures. Biomaterials 2001, 22 (10), 1077–84.

43. Wang, S. S.; Becerra-Arteaga, A.; Good, T. A., Development of a novel diffusion-based method to estimate the size of the aggregated Abeta species responsible for neurotoxicity. Biotechnol Bioeng 2002, 80 (1), 50–9.

44. Laurent, T. C.; Fraser, J. R. E., Hyaluronan. FASEB J 1992, 6, 2397–2404.

45. Ruoslahti, E., Brain extracellular matrix. Glycobiology 1996, 6 (5), 489–492.

46. Suri, S.; Schmidt, C. E., Cell-laden hydrogel constructs of hyaluronic acid, collagen, and laminin for neural tissue engineering. Tissue Eng Part A 2010, 16 (5), 1703–16.

47. Xu, X.; Jha, A. K.; Harrington, D. A.; Farach-Carson, M. C.; Jia, X., Hyaluronic Acid-Based Hydrogels: from a Natural Polysaccharide to Complex Networks. Soft Matter 2012, 8 (12), 3280–3294.

48. Aljohani, W.; Ullah, M. W.; Zhang, X.; Yang, G., Bioprinting and its applications in tissue engineering and regenerative medicine. Int J Biol Macromol 2018, 107 (Pt A), 261–275.

49. Feng, Q.; Zhu, M.; Wei, K.; Bian, L., Cell-mediated degradation regulates human mesenchymal stem cell chondrogenesis and hypertrophy in MMP-sensitive hyaluronic acid hydrogels. PLoS One 2014, 9 (6), e99587.

50. Khetan, S.; Guvendiren, M.; Legant, W. R.; Cohen, D. M.; Chen, C. S.; Burdick, J. A., Degradation-mediated cellular traction directs stem cell fate in covalently crosslinked three-dimensional hydrogels. Nat Mater 2013, 12 (5), 458–65.

51. Fisher, S. A.; Anandakumaran, P. N.; Owen, S. C.; Shoichet, M. S., Tuning the Microenvironment: Click-Crosslinked Hyaluronic Acid-Based Hydrogels Provide a Platform for Studying Breast Cancer Cell Invasion. Adv Funct Mater 2015, 25 (46), 7163–7172.

52. Cui, F. Z.; Tian, W. M.; Hou, S. P.; Xu, Q. Y.; Lee, I. S., Hyaluronic acid hydrogel immobilized with RGD peptides for brain tissue engineering. J Mater Sci Mater Med 2006, 17 (12), 1393–401.

53. Leach, J. B.; Bivens, K. A.; Patrick, C. W., Jr.; Schmidt, C. E., Photocrosslinked hyaluronic acid hydrogels: natural, biodegradable tissue engineering scaffolds. Biotechnol Bioeng 2003, 82 (5), 578–89.

54. Marklein, R. A.; Burdick, J. A., Spatially controlled hydrogel mechanics to modulate stem cell interactions. Soft Matter 2010, 6 (1), 136–143.

55. Khan, R.; Mahendhiran, B.; Aroulmoji, V., Chemistry of hyaluronic acid and its significance in drug delivery strategies: A review. Int J Pharm Sci Res. 2013, 4 (9), 3699–3710.

56. Shu, X. Z.; Liu, Y.; Luo, Y.; Roberts, M. C.; Prestwich, G. D., Disulfide Cross-Linked Hyaluronan Hydrogels. Biomacromolecules 2002, 3 (6), 1304–1311.

57. Nimmo, C. M.; Owen, S. C.; Shoichet, M. S., Diels-Alder Click cross-linked hyaluronic acid hydrogels for tissue engineering. Biomacromolecules 2011, 12 (3), 824–30.

58. Wieland, J. A.; Houchin-Ray, T. L.; Shea, L. D., Non-viral vector delivery from PEG-hyaluronic acid hydrogels. J Control Release 2007, 120 (3), 233–41.

59. Jin, R.; Dijkstra, P. J.; Feijen, J., Rapid gelation of injectable hydrogels based on hyaluronic acid and poly(ethylene glycol) via Michael-type addition. J Control Release 2010, 148 (1), e41–3.

60. Cruise, G. M.; Scharp, D. S.; Hubbell, J. A., Characterization of permeability and network structure of interfacially photopolymerized poly(ethylene glycol) diacrylate hydrogels. Biomaterials 1998, 19 (14), 1287–1294.

61. Edgar, L.; McNamara, K.; Wong, T.; Tamburrini, R.; Katari, R.; Orlando, G., Heterogeneity of Scaffold Biomaterials in Tissue Engineering. Materials (Basel) 2016, 9 (5).

62. Zustiak, S. P.; Leach, J. B., Hydrolytically degradable poly(ethylene glycol) hydrogel scaffolds with tunable degradation and mechanical properties. Biomacromolecules 2010, 11 (5), 1348–57.

63. Phelps, E. A.; Enemchukwu, N. O.; Fiore, V. F.; Sy, J. C.; Murthy, N.; Sulchek, T. A.; Barker, T. H.; Garcia, A. J., Maleimide cross-linked bioactive PEG hydrogel exhibits improved reaction kinetics and cross-linking for cell encapsulation and in situ delivery. Adv Mater 2012, 24 (1), 64–70, 2.

64. Waters, J., The concentration of soluble extracellular amyloid-beta protein in acute brain slices from CRND8 mice. PLoS One 2010, 5 (12), e15709.

65. Sengupta, P.; Garai, K.; Balaji, J.; Periasamy, N.; Maiti, S., Measuring Size Distribution in Highly Heterogeneous Systems with Fluorescence Correlation Spectroscopy. Biophys J 2003, 84 (3), 1977–1984.

66. Duarte Campos, D. F.; Bonnin Marquez, A.; O’Seanain, C.; Fischer, H.; Blaeser, A.; Vogt, M.; Corallo, D.; Aveic, S., Exploring Cancer Cell Behavior In Vitro in Three-Dimensional Multicellular Bioprintable Collagen-Based Hydrogels. Cancers (Basel) 2019, 11 (2).

67. Velez, D. O.; Tsui, B.; Goshia, T.; Chute, C. L.; Han, A.; Carter, H.; Fraley, S. I., 3D collagen architecture induces a conserved migratory and transcriptional response linked to vasculogenic mimicry. Nat Commun 2017, 8 (1), 1651.

68. Ahn, J.; Ahn, J. H.; Yoon, S.; Nam, Y. S.; Son, M. Y.; Oh, J. H., Human three-dimensional in vitro model of hepatic zonation to predict zonal hepatotoxicity. J Biol Eng 2019, 13, 22.

69. Bahcecioglu, G.; Hasirci, N.; Bilgen, B.; Hasirci, V., A 3D printed PCL/hydrogel construct with zone-specific biochemical composition mimicking that of the meniscus. Biofabrication 2019, 11 (2), 025002.

70. Manning, K. L.; Thomson, A. H.; Morgan, J. R., Funnel-Guided Positioning of Multicellular Microtissues to Build Macrotissues. Tissue Eng Part C Methods 2018, 24 (10), 557–565.

71. Wu, S.; Xu, R.; Duan, B.; Jiang, P., Three-Dimensional Hyaluronic Acid Hydrogel-Based Models for In Vitro Human iPSC-Derived NPC Culture and Differentiation. J Mater Chem B 2017, 5 (21), 3870–3878.

72. Xiao, W.; Ehsanipour, A.; Sohrabi, A.; Seidlits, S. K., Hyaluronic-Acid Based Hydrogels for 3-Dimensional Culture of Patient-Derived Glioblastoma Cells. J Vis Exp 2018, (138).

73. Sharma, P. K.; Taneja, S.; Singh, Y., Hydrazone-Linkage-Based Self-Healing and Injectable Xanthan-Poly(ethylene glycol) Hydrogels for Controlled Drug Release and 3D Cell Culture. ACS Appl Mater Interfaces 2018, 10 (37), 30936–30945.

74. Bicker, J.; Alves, G.; Fortuna, A.; Falcao, A., Blood-brain barrier models and their relevance for a successful development of CNS drug delivery systems: a review. Eur J Pharm Biopharm 2014, 87 (3), 409–32.

75. Hortschansky, P.; Schroeckh, V.; Christopeit, T.; Zandomeneghi, G.; Fandrich, M., The aggregation kinetics of Alzheimer’s beta-amyloid peptide is controlled by stochastic nucleation. Protein Sci 2005, 14 (7), 1753–9.

76. Streets, A. M.; Sourigues, Y.; Kopito, R. R.; Melki, R.; Quake, S. R., Simultaneous measurement of amyloid fibril formation by dynamic light scattering and fluorescence reveals complex aggregation kinetics. PLoS One 2013, 8 (1), e54541.

77. Weber, P. A.; Chang, H. C.; Spaeth, K. E.; Nitsche, J. M.; Nicholson, B. J., The permeability of gap junction channels to probes of different size is dependent on connexin composition and permeant-pore affinities. Biophys J 2004, 87 (2), 958–73.

78. Sengupta, P.; Garai, K.; Balaji, J.; Periasamy, N.; Maiti, S., Measuring Size Distribution in Highly Heterogeneous Systems with Fluorescence Correlation Spectroscopy. Biophysical Journal 2003, 84 (3), 1977–1984.

79. Amaro, M.; Wellbrock, T.; Birch, D. J. S.; Rolinski, O. J., Inhibition of beta-amyloid aggregation by fluorescent dye labels. Applied Physics Letters 2014, 104 (6), 063704.

80. Wu, C.; Lei, H.; Wang, Z.; Zhang, W.; Duan, Y., Phenol red interacts with the protofibril-like oligomers of an amyloidogenic hexapeptide NFGAIL through both hydrophobic and aromatic contacts. Biophys J 2006, 91 (10), 3664–72.

81. Reyes Barcelo, A. A.; Gonzalez-Velasquez, F. J.; Moss, M. A., Soluble aggregates of the amyloid-beta peptide are trapped by serum albumin to enhance amyloid-beta activation of endothelial cells. J Biol Eng 2009, 3, 5.

