## Supplemental Figure 1 for "Impact of four common hydrogels on amyloid-β (Aβ) aggregation and cytotoxicity: Implications for 3D models of Alzheimer’s disease"

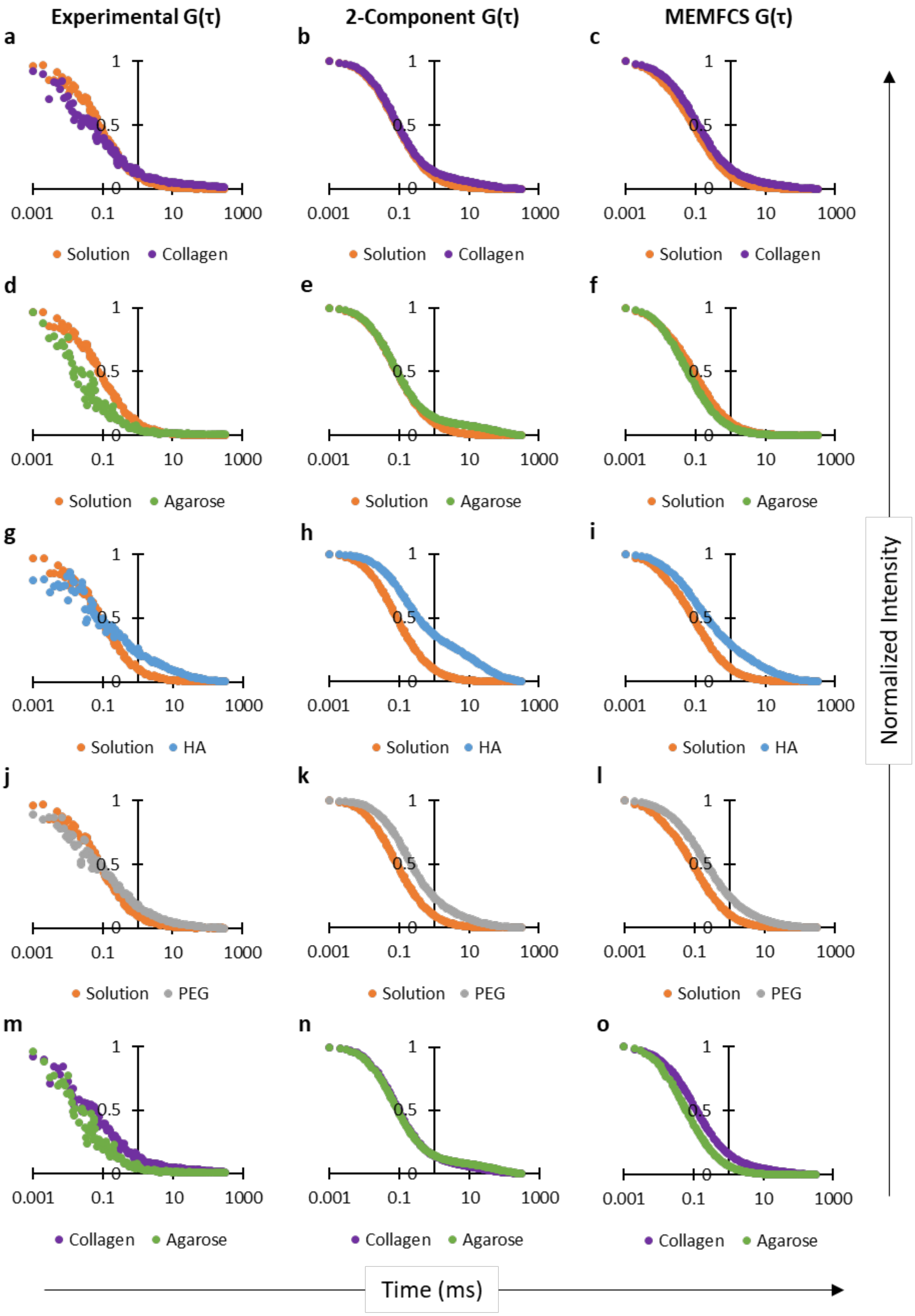

Experimental G(τ)
2-Component G(τ)
MEMFCS G(τ)
a
b
c
d
e
f
g
h
i
j
k
l
m
n
o
1
0.5
0
0.001
0.1
10
1000
Solution
Collagen
Agarose
HA
PEG
Normalized Intensity
Time (ms)

***Supplementary Figure 1*** *G($\tau$) comparison curves of raw experimental data, calculated 2-component model fits, and calculated MEMFCS model fits (Part 1).*

*Experimental comparisons are in the first column 2-component model comparisons are in the middle column, and MEMFCS model comparisons are in the third column. Solution data is in orange, collagen data is in purple, agarose data is in green, HA data is in blue, and PEG data is in grey. Row **a-c** compares solution and collagen. Row **d-f** compares solution and agarose. Row **g-I** compares solution and HA. Row **j-l** compares solution and PEG. Row **m-o** compares collagen and agarose. The p-values in Supplementary Table 1 correspond to these figure comparisons.*
