## Supplemental Figure 2 for "Impact of four common hydrogels on amyloid-β (Aβ) aggregation and cytotoxicity: Implications for 3D models of Alzheimer’s disease"

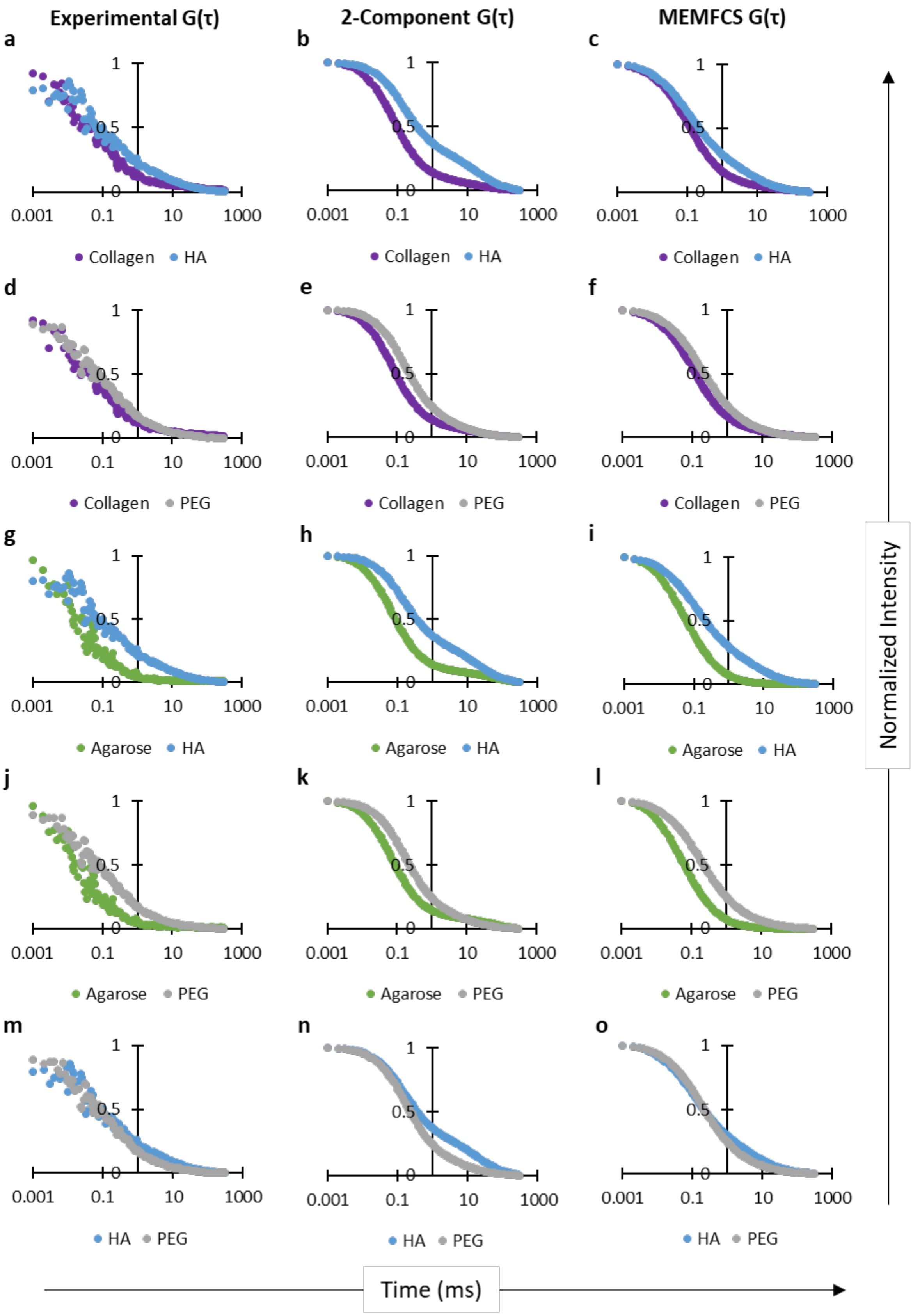
Experimental G(τ)
2-Component G(τ)
MEMFCS G(τ)
a
b
c
d
e
f
g
h
i
j
k
l
m
n
o
1
0.5
0
0.001
0.1
10
1000
Collagen
HA
PEG
Agarose
Normalized Intensity
Time (ms)

1

***Supplementary Figure 1*** *G($\tau$) comparison curves of raw experimental data, calculated 2-component model fits, and calculated MEMFCS model fits (Part 2).*

*Experimental comparisons are in the first column 2-component model comparisons are in the middle column, and MEMFCS model comparisons are in the third column. Solution data is in orange, collagen data is in purple, agarose data is in green, HA data is in blue, and PEG data is in grey. Row **a-c** compares collagen and HA. Row **d-f** compares collagen and PEG. Row **g-I** compares agarose and HA. Row **j-l** compares agarose and PEG. Row **m-o** compares HA and PEG. The p-values in Supplementary Table 1 correspond to these figure comparisons.*
