## Supplementary Table 1 for "Impact of four common hydrogels on amyloid-β (Aβ) aggregation and cytotoxicity: Implications for 3D models of Alzheimer’s disease"

| | | | Experimental G(τ) | 2-Component G(τ) | MEMFCS G(τ) |
|---|---|---|---|---|---|
| Solution | - | Collagen | **<0.0001** | **0.0003** | **0.0004** |
| Solution | - | Agarose | **0.0004** | **<0.0001** | 0.8994 |
| Solution | - | HA | **<0.0001** | **<0.0001** | **<0.0001** |
| Solution | - | PEG | **0.0056** | **0.0004** | **0.001** |
| Collagen | - | Agarose | **<0.0001** | 0.3014 | **<0.0001** |
| Collagen | - | HA | **0.0171** | **<0.0001** | **0.0463** |
| Collagen | - | PEG | 0.112* | 0.1888 | 0.5436 |
| Agarose | - | HA | **<0.0001** | **<0.0001** | **<0.0001** |
| Agarose | - | PEG | **0.0002** | 0.1888 | **<0.0001** |
| HA | - | PEG | 0.1464* | **0.0025** | 0.454 |

***Supplementary Table 1*** *Comparative KS 2-sample test of G(τ) curves.*

*With 95% confidence, p-values <0.05 were significantly different. The false discovery rate (FDR) was set at 5% using the two-stage step-up method of Benjamini, Krieger and Yekutieli. For the experimental G(τ), the FDR test found all the samples significant, shown with an "*". For the 2-component model and the MEMFCS model, the FDR test did not change the significance of any comparisons.*
